# From ancestral patterns of variation to the human condition: Leveraging disease phenotype associations and gene expression changes

**DOI:** 10.64898/2026.11.22.768481

**Authors:** Elif M. Yildiz, Gonzalo Barrio, Yulia Yakubenkova, Cedric Boeckx

## Abstract

The shift in perspective to characterize the genetic basis of the human condition beyond fully fixed mutations opens many possibilities to scrutinize the nature of high-frequency derived variants in modern human genomes through the effect of the minor alleles, either already present in extant *Pan* relatives (“basal”), or found across most genomes of extinct hominins and likely introgressed into *sapiens* genomes. Here we present the results of intersecting a comprehensive catalog of such loci with the ClinVar and gnomAD databases, which provide associations with disease phenotypes, valuable to infer functional impacts. Recurrent disease associations highlight terms with sensorimotor processing and ataxia. Examining the impact of these variants in the GTEx database reveals striking differences between “basal” and “derived” patterns of variation: In contrast to minor alleles traceable to the Pan lineage (“basal variation”), minor alleles that are derived in extinct hominin genomes are found in highly constrained genomic regions in the GnomAD database, and are in general depleted of brain-specific effects in the GTEx database. Our Gtex analysis also highlights the cerebellum as the tissue with the most salient differences between major and minor (basal) alleles, pointing to the relevance of this brain region for understanding the evolution of our lineage. This finding is further reinforced by a Gene Ontology pathway analysis, which brings out metabolic terms, some specific to the cerebellum and glial cell types.

## 1. Introduction

Over a decade of work on high-coverage genomes from extinct late hominins (hereafter, Neanderthals and Denisovans) and how these differ systematically from contemporary human genomes has shifted the focus away from “fully exclusive” variants (derived alleles that are fully fixed in one lineage) to variants exhibiting a non-exclusive but nonetheless highly divergent pattern: derived, and found at very high frequency (>90%) in one lineage (say, *sapiens*), but very rare, or virtually absent in the other lineage (say, Neanderthal).

Single nucleotide polymorphisms of this kind have been used to rethink the “human condition” from a molecular perspective (Pääbo, 2014; Kuhlwilm & Boeckx, 2019; Zeberg et al., 2024). The initial, natural hypothesis of a set of exclusively derived, changes universally shared in one population but absent in a closely related population has gradually made way to a view where lineage-specific traits are thought to be “the consequence of cumulative changes” that form a “network” in which no single change needs to be fixed in all the individuals (Kuhlwilm & Boeckx, 2019). This “combinatorial view of what constitutes, say, a modern human” (as opposed to a Neanderthal, a Denisovan, or even a representative of an early *sapiens* group) preserves the idea that “some individual genetic changes may be more functionally important than others” but recognizes that each individual of the relevant lineage needs to carry most but not necessarily all of these genetic features (Zeberg et al., 2024). The main reason for this shift is that, as more and more genomes are sequenced, evidence of instances of (bidirectional) gene flow is increasing. As a result, the number of completely fixed derived variants decreases rapidly.

In light of this shift of perspective towards a pluralistic, combinatorial view of the human condition (“human conditions”, Caporale et al., 2025), we set out to examine patterns of “highly” divergent polymorphisms in order to gain further insights into the functional consequences of changes derived in one, but not the other late hominin lineages. We focused specifically on polymorphic sites annotated in the ClinVar database (https://www.ncbi.nlm.nih.gov/clinvar/), which aggregates information about genomic variation in contemporary humans and its relationship to human health. Crucially, our intention was not to exploit such associations with diseases to make claims about the prevalence of such diseases in extinct hominin lineages, as such associations or risk scores are known not to transfer well across diverse populations, even within a single species (Ding et al. 2023). Instead, our aim was to use these associations and what we know about associated diseases to ask questions about the more proximate phenotypic differences between the highly divergent major and minor alleles, such as cellular consequences or selective anatomical impact (e.g., do they cluster around a specific tissue?). That is to say, our aim was to exploit the disease association---especially so, if the disease affects some tissues more than others---as an additional source of information guiding the functional interpretation of specific polymorphic sites, hypothesizing that whereas disease phenotypes do not transfer across populations, locus of divergent expression or “molecular phenotypes” (Brand et al., 2022) are more likely to do so. With this aim in mind, we examined the impact of disease-associated variants with tissue-specific information provided in the Gtex Portal.

The “highly” divergent polymorphisms we decided to focus on all separate sapiens from Neanderthals and Denisovans (following Kuhlwilm & Boeckx, 2019, the major allele in sapiens is found in >90% of the global contemporary human population, and the minor allele in us is most common across other hominin genomes), but come in two varieties (Figure 1A): one where the minor sapiens allele is the major allele not only in Neanderthals and Denisovans but also in closely related extant primate species like chimpanzees, and another where the major allele in Neanderthals and Denisovans constitutes a derived allele not present in chimpanzees. We call this first variety type “basal” to indicate that this type of variation likely corresponds to an instance of a very “old” pattern of polymorphism, where one ancestral variant was progressively replaced by a newer variant derived in the *sapiens* lineage. We call the second variety type “(Neanderthal/Denisovan)-derived”. For this second type, it is likely that the minor allele in the *sapiens* branch was introduced as a result of gene flow among hominins, of which several instances are now well attested (Huang et al., 2025; Ongaro & Huerta-Sanchez, 2024; Gokcumen, 2020; Skov et al., 2020). But it is also possible that the minor allele on the *sapiens* branch may be the result of incomplete lineage sorting and was present in the last common ancestor among late hominins. The lack of genomes from that last common ancestor makes it difficult to fully disentangle these two options. From the geographical distribution of the derived minor allele examined here (much rarer in Africa than in Europe or Asia), we suspect that introgression was the major factor, but we do not seek to make any strong claim on this point in this study. As far as the basal variety is concerned, we also recognize the possibility that the minor allele in *sapiens* may have been lost early in our lineage only to be reintroduced back via introgression from Neanderthal/Denisovan populations (Rinker et al., 2020), based on published work on introgressed variants, but we think that this was a minor factor underlying basal patterns of variation examined here.

**Figure 1.**
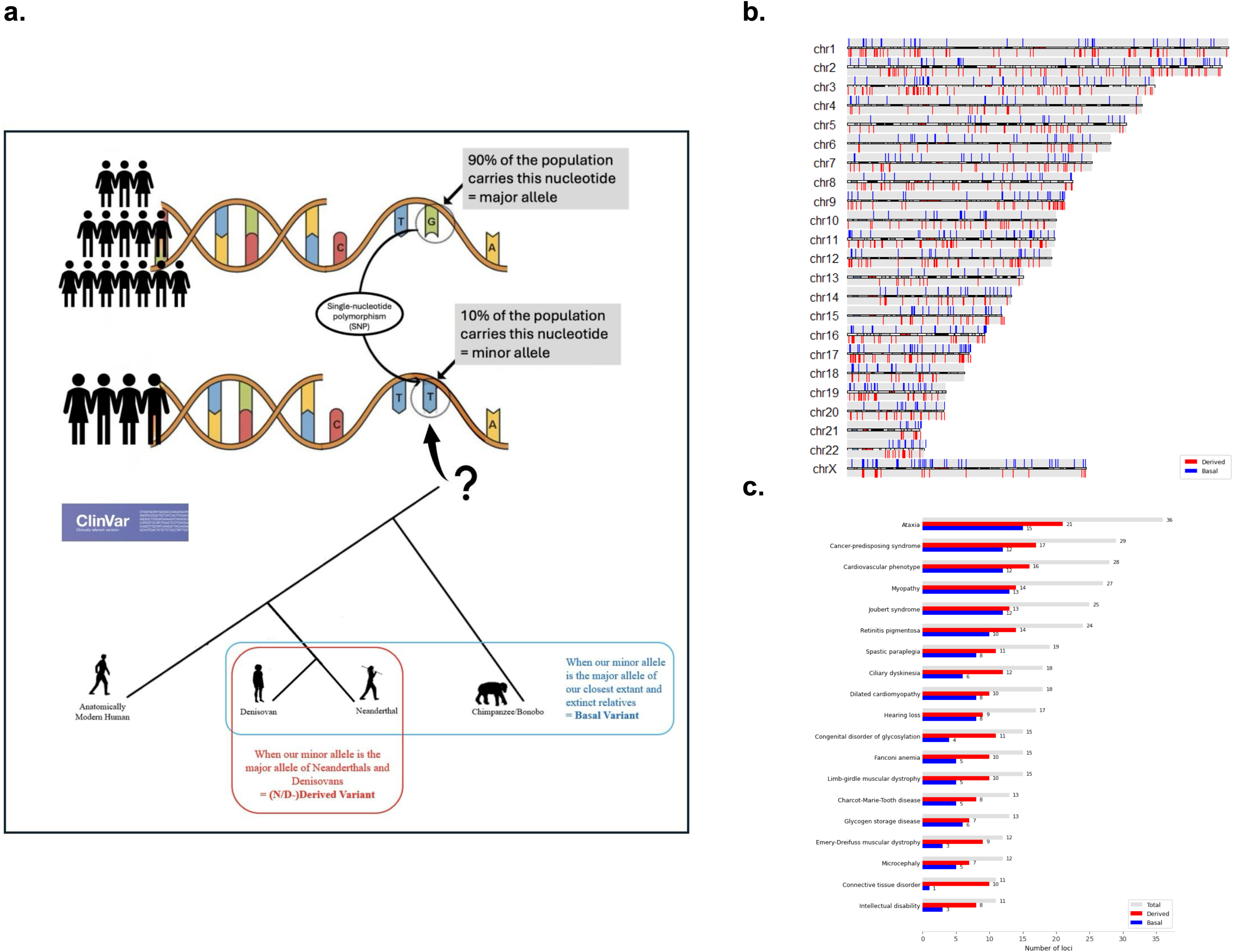
Characterization of alleles under investigation. a. Schematic representation of the two classes of variants investigated: basal variants, for which the minor allele in *Homo sapiens* is also found in Neanderthals, Denisovans and other extant primates; and (N/D)-derived variants, for which the minor allele in *Homo sapiens* corresponds to the major allele in Neanderthals and Denisovans but is absent from other great apes. b. Distribution of basal and (N/D)-derived variants across the human karyotype. c. Most frequent disease and phenotype associations in ClinVar across the total dataset and the basal and (N/D)-derived subsets.

## 2. Results

### 2.1. Overall distribution of ClinVar variants

We filtered 136,435 high-frequency variants catalogued by Kuhlwilm and Boeckx (2019) through the ClinVar database, which resulted in the identification of a total of 711 basal SNPs and 957 derived SNPs (Table S1). All the variants are approximately equally distributed along the karyotype (Figure 1b), with no remarkable differences between the basal and derived SNPs overall.

Out of the 1,668 total SNPs covered by ClinVar, 688 (41.25%) did not return specific information about the disease associated with the variant. The remaining 980 were associated with 578 different diseases (Table S2). The 432 basal variants with defined associations pointed to 322 distinct pathologies, 185 (57.28%) of which were uniquely found in this subset, while the 548 derived SNPs with known labels spanned 394 diseases, 256 (64.97%) being exclusive to this list. Significantly, 138 types appeared in both sets and constituted the majority of the highly prevalent categories. Most diseases (313), on the other hand, are associated with only 1 variant in either one of the lists and do not allow for systematic analysis.

We therefore focused on the recurrent disease associations across both basal and derived categories, which revealed a set of disease categories represented across multiple variants, spread across the genome and targeting distinct genes (Figure 1c). Ataxia was the most frequent association term, driven largely by non-overlapping basal and derived loci. Other recurrent associations included sensory conditions such as myopathy, retinitis pigmentosa, and hearing loss; ciliopathies such as Joubert syndrome and ciliary dyskinesia; specific metabolic disorders, including glycosylation and glycogen storage disorders; and muscular dystrophy.

To probe whether such recurrent disease associations may be driven by associations with long genes (known to enrich for Central Nervous System associations, Zylka et al., 2015), we sought to annotate basal/derived SNPs with genes that are likely to impact them (Table S3). Gene prediction was conducted with different tools to obtain a set of credible candidate genes in our dataset. The Genomic Regions Enrichment of Annotations Tool (GREAT) proposed a total of 1,856 genes possibly associated with the high frequency variants that was narrowed down by comparing it to the predictions provided by the Genome Aggregation Database (gnomAD; https://gnomad.broadinstitute.org/). Selecting only genes that were predicted both in GREAT and gnomAD for a given variant resulted in a set of 957 candidate genes (0.57% of the initial dataset), of which 450 were associated with the basal variants and 636 with the derived ones, with some overlap between the genes in two sets of variants.

We asked whether there a difference in the length of genes associated with the ten most frequent disease categories compared with randomly selected control genes and found no significant difference in either basal (p = 0.244) or derived (p = 0.481) sets (Table S3).

### 2.2. GTEx matches

To further understand the impact of basal/derived SNPs, we searched the GTEx database (https://gtexportal.org/home/) to determine if any of our SNPs were attested expression and splicing Quantitative Trait Loci (e-s/QTLs), aiming to detect any tissue enrichment. We consulted both GTEx version 8 and the latest, version 10. While results are broadly similar (Table S14), we note a slight 1.2% increase of eQTL results but a more significant 81.8% increase of sQTL results with GTEx 10. The results presented in the main text correspond to GTEx 10 (Table S4).

In GTEx 10, our dataset of 1,668 variants contained 340 significant eQTLs (20.38%) and 245 significant sQTLs (14.69%). In the basal subset of 711 variants, 293 were identified as eQTLs (41.21%), 216 as sQTLs (30.37%) and 184 as both (25.88%). In the derived subset of 957 variants, on the other hand, only 47 were reported as eQTLs (4.91%), 29 as sQLTs (3.03%) and 17 as both (1.78%). This uncovers a significant depletion of QTL results in the derived set. This is especially noteworthy in light of the fact that the initial derived SNP set is bigger than the basal one. We considered several reasons for this salient asymmetry and concluded that the most likely cause is allele frequency (Figure 2a). Although both basal and derived datasets conform to the <10% frequency for the minor allele in the current global human population, their MAF distributions differ significantly. Basal variants have a mean MAF of 0.0361, compared with 0.0017 for derived variants (Welch’s two-sample t-test: *t* = 414.39, *p* < 2.2 × 10⁻¹⁶). Thus, basal variants tend to be considerably more frequent in present-day human populations than derived variants. Being rarer, N/D-derived minor alleles may be less well represented in databases like GTEx. A prominent reason for this frequency difference may of course be the result of geographically limited introgression patterns: only a portion of the world’s population exhibits non-negligible traces of ancient gene flow.

**Figure 2.**
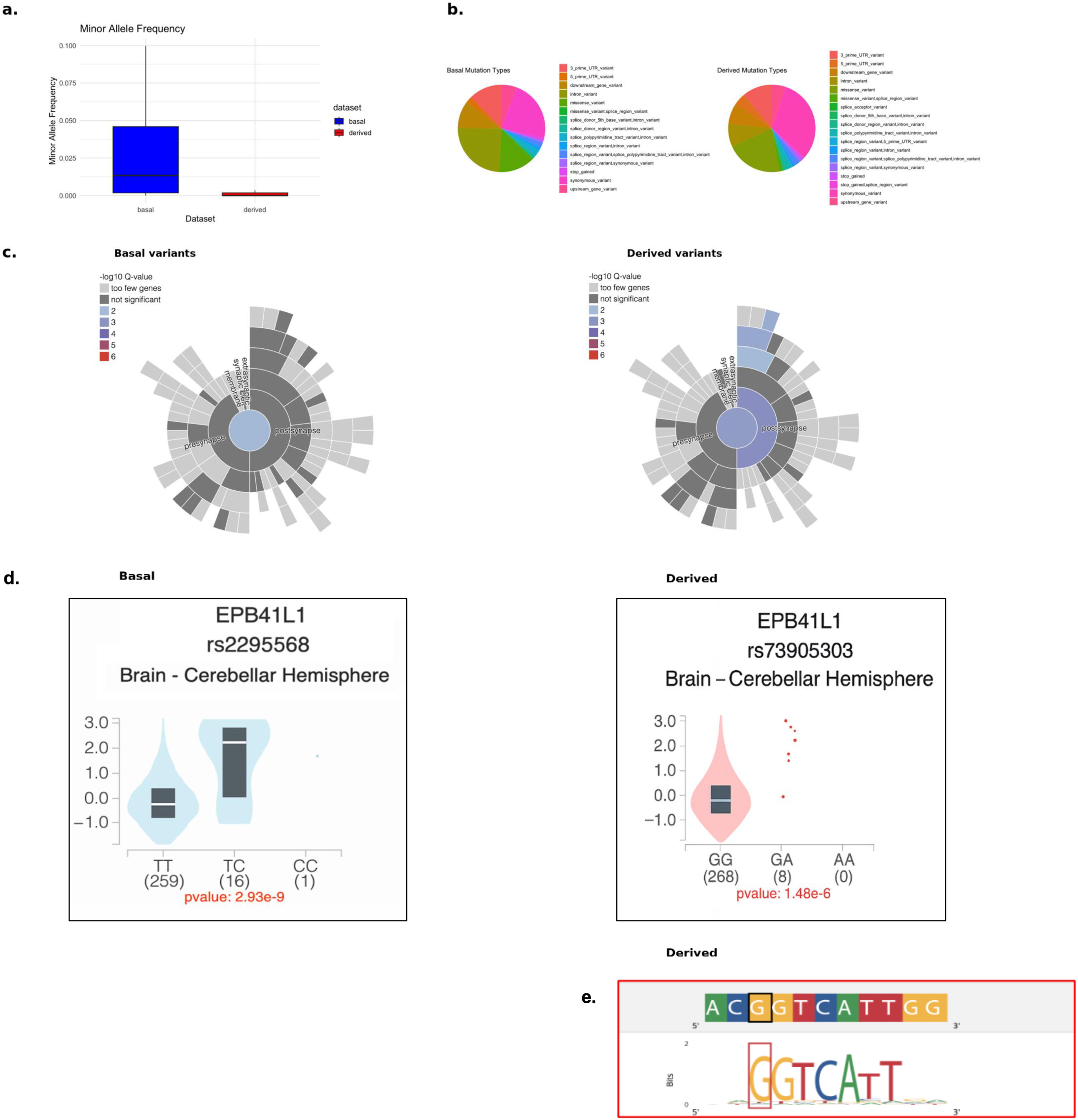
Basal/Derived variant asymmetries. a. Minor allele frequency differences between basal and derived variants. b. Mutation type asymmetries between basal and derived variants. c. SYNGO-enrichment differences between basal and derived variants. d. Genotype-dependent EPB41L1 expression associated with the basal (rs2295568) and (N/D)-derived (rs73905303) minor alleles in the cerebellar hemisphere, illustrating differences in their effects on gene expression. e. Sequence logo illustrating the predicted transcription-factor binding motif surrounding the (N/D)-derived minor allele.

However, other factors may also be at play regarding the asymmetry uncovered between basal and derived sets (Figure 2b). Both basal and derived SNP lists contain an equal amount of minor alleles labeled “pathogenic” in ClinVar, but the derived set contains a higher number of variants labeled “of uncertain significance”. We therefore examined genetic constraint at both the gene and genomic levels, using GnomAD (see Methods; Table S5). At the gene level, constraint scores were broadly comparable between the two sets, with no significant differences observed for synonymous variants (basal mean = −0.47, derived = −0.29; p = 0.182), missense variants (1.02 vs 1.08; p = 0.996), or pLoF-associated genes (LOEUF: 0.84 vs 0.87; p = 0.591). At the genomic level, Gnocchi constraint scores were higher for basal variants than derived variants (mean = 0.82 vs 0.64), although this difference was not significant (Wilcoxon p = 0.138; Welch’s t-test p = 0.063). Using a threshold of constraint_z ≥ 2, 25.5% of basal variants and 23.3% of derived variants fell into the high-constraint category, with no significant difference in enrichment between the groups (Fisher’s exact test, p = 0.313). Having said this, the standard Ensembl Variant Effect Predictor (VEP), used to annotate the genomic consequences of variants based on their location relative to genes and transcripts, revealed differences in mutation-impact distributions between basal and derived variants. Using VEP-defined impact categories (MODIFIER, LOW, MODERATE and HIGH), derived variants were significantly more likely to fall into higher-impact categories than basal variants (ordinal logistic regression: β = 0.70, SE = 0.07, z = 10.51, p < 0.001, OR ≈ 2.01; see Methods and Table S6).

We further assessed the predicted functional impact of individual variants using CADD PHRED scores (Table S7). Derived eQTL variants showed the highest mean PHRED scores overall (∼8.16), and the largest effect was observed between derived eQTL and derived sQTL variants (Cohen’s d = 0.82), indicating a large difference in predicted functional impact between these groups. Medium-to-large effect sizes were also observed when comparing derived eQTL variants with the basal groups (d ≈ 0.60–0.65). However, none of the pairwise comparisons remained statistically significant following Bonferroni correction, likely reflecting the small sample sizes of the derived eQTL (n = 19) and derived sQTL (n = 12) groups together with the conservativeness of the correction.

We close this section by observing two additional differences between basal and derived sets. Using GREAT-predicted gene associations for all basal and all Neanderthal/Denisovan-derived SNPs, which revelead the presence of quite a few autism-related (SFARI) genes (Table S12), we observed differences in brain-related enrichment patterns between the two groups, as assessed using the SynGO platform (https://www.syngoportal.org/): The basal and derived SNP sets showed differing enrichment patterns across synaptic compartments (Figure 2c).

In addition, we observe that in a handful of cases where we find both a basal and derived SNP significantly impacting the expression of the same gene in the same tissue, as reported in GTEx, the derived SNP appears more disruptive (affecting a Transcription Binding site), as is the case for rsIDs rs2295568 and rs73905303, according to both the Regulome (https://www.regulomedb.org/) and the UCSC platforms (https://genome.ucsc.edu/) (Figure 2d-e). We could not detect the opposite pattern (basal more disruptive) in our dataset.

### 2.3. Tissue specificity

To further understand the distribution of QTLs across tissues, we first assessed whether the observed tissue distribution differed from that expected based on GTEx sampling proportions. The tissue distributions differed significantly from GTEx sampling proportions for basal eQTLs (pFDR = 2 × 10⁻⁶) and basal sQTLs (pFDR = 2 × 10⁻⁶), as well as derived sQTLs (pFDR = 4.88 × 10⁻⁴), whereas derived eQTLs did not differ significantly from the expected distribution (pFDR = 0.459) (Table S4). We note that brain tissues systematically exhibit more extreme (regardless of whether the variant up-regulates or down-regulates the gene) normalised expression size (NES) than non-brain tissues as reflected in the broader tail of the brain NES distribution (Figure 3a).

**Figure 3.**
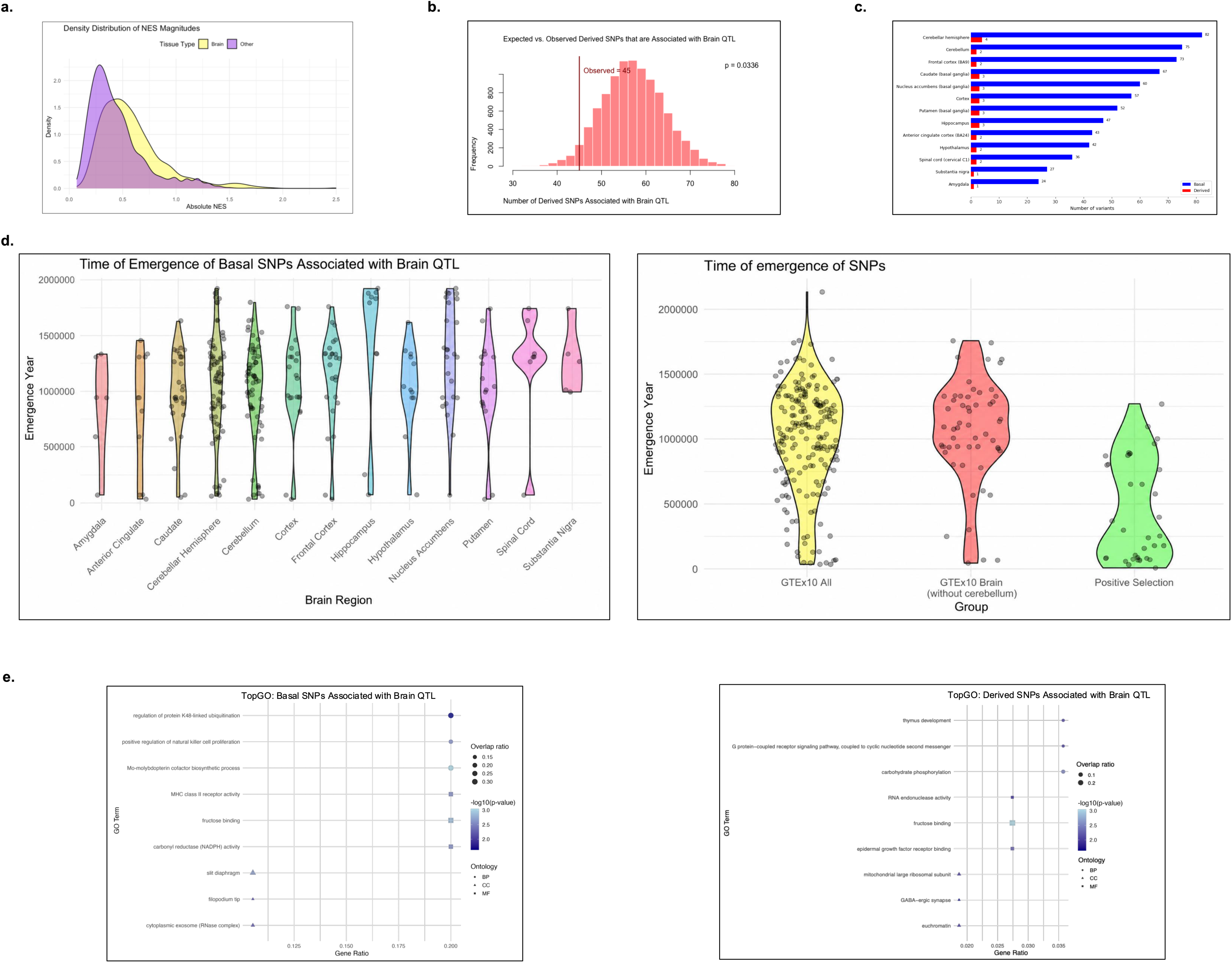
QTL results. a. Density distributions of absolute normalised effect sizes (NES) in brain and non-brain tissues. b. Permutation-based assessment of the depletion of brain QTL-associated variants among (N/D)-derived variants, showing the expected distribution under the null hypothesis and the observed number of variants. c. Distribution of basal and (N/D)-derived QTL-associated variants across individual brain tissues. d. Estimated times of emergence of basal variants associated with QTL results across individual brain tissues (left), and comparison of emergence-time distributions across all GTEx QTL-associated variants, brain QTL-associated variants (excluding the cerebellum), and QTL-associated variants overlapping regions with signals of positive selection (right). e. TopGO enrichment terms for basal variants associated with brain QTLs across all brain tissues (left) and cerebellum/cerebellar hemisphere QTLs (right).

We then specifically tested whether the representation of brain-associated QTLs differed between the basal and derived sets. A permutation analysis preserving the brain/non-brain status of SNPs showed that the low number of brain QTLs in the derived group was unlikely under the null distribution (p = 0.036), supporting a significant under-representation of brain QTLs among derived variants (Figure 3b).

For the basal set, among brain tissues, we find that the cerebellum and cerebellar hemisphere occur most frequently, even after controlling for SNPs that have adjacent genomic coordinates and are associated with the same gene (Figure 3c). This replicates an earlier finding examining derived, *Homo sapiens*-specific high-frequency alleles that are associated with differential gene expression across 15 brain structures derived from GTEx 7 (Andirkó & Boeckx, 2022), and is in line with other studies pointing to a variety of differences in the cerebellum among hominins (Gunz et al., 2019; McCoy et al., 2017; Buisan et al., 2022; Andirkó et al., 2022), associated with marked changes in the cerebellum late in our lineage (Neubauer et al., 2018). For N/D-derived SNPs, where the numbers are considerably smaller, the caudate is impacted as much as the cerebellum, consistent with claims in Mafessoni et al. (2020), Gunz et al. (2019) and McCoy et al. (2017).

In light of this, we plotted the times of emergence for basal variants found in GTEx, using the open repository of genetic variant age estimations (Albers & McVean, 2020; Andirkó et al., 2022) (Figure 3d). Compared to the overall profile found for all derived high-frequency gene variants reported in Andirkó et al. (2022) (bimodal distribution), we find a higher concentration of older variants (i.e., older than 500kya) among the basal variants under examination (Table S9). Among the basal variants found in GTEx impacting brain tissues, SNPs impacting the cerebellum/cerebellar hemisphere contribute the most to the group of variants that are dated between 200kya and present-day. Variants impacting other brain tissues tend to be of much older origin.

We performed functional enrichment analyses on all brain e/s-genes and on cerebellum/cerebellar hemisphere e/s-Genes for the basal dataset using a Gene Ontology pathway analysis (Table S10). e/s-Genes associated with the brain (all regions) are enriched for general metabolic processes and enzymatic activity. Immune regulation processes are also over-represented (Figure 3e). On the other hand, e/s-genes associated with the cerebellum/cerebellar hemisphere enrich more specific neuron-related (GABAergic synapse) and signalling processes (G protein-coupled receptor signalling pathway), in addition to metabolic terms.

The specific metabolic terms (carbohydrate metabolism, glycolysis, fructose metabolism) are known to be associated with glucose metabolism and are more closely associated with some cell types, such as astrocytes and other glia, at adult stages (matching the GTEx database). This led us to probe the cellular specificity of e/s-genes in our dataset against a comprehensive resource like Siletti et al. (2023) (Table S11). Higher expression of basal e/s-genes was found in a variety of cells associated with the epithelium and vasculature, although the cell number of these populations were small. By contrast, the cluster of “immature astrocyte” cells was very high and contained more e/s-genes than canonical neuronal cell types like excitatory or inhibitory neurons.

Prominent genes associated with specific metabolic functions among our datasets include *FCSK*; *KHK*; *PFKM*; *HK2*; *ACADVL*; *FOXK1*; *GYS1*; *SLC2A1*; *HINT1*. Many of these feature in the literature on astrocyte energetics. Quite a few of them are also known to play a role in embryonic brain development. The current GTEx database does not include fetal data, but we examined a study (Wen et al., 2024) providing QTL data for prenatal cortical development and identified several matching QTLs (those enriching for metabolic terms like carbohydrate metabolism). Of note, in the context of prenatal development, several of our SNPs fall in sequences that show active regulatory activity and drive differential expression between *Homo* groups in a massively parallel reporter assay in embryonic stem cells, neural progenitor cells, and bone osteoblasts (Weiss et al., 2021). Two of these instances are predicted to significantly impact the expression of well-studied autism candidate genes: *CACNA1C*, long associated with schizophrenia and Timothy Syndrome, and *SHANK2*, a high confidence autism gene. No GTEx information was available for the relevant SNPs, though. The only highly confident SFARI gene present as an e-gene in our dataset is *CDKL5*.

Consistent with the presence of ciliopathies among disease enrichments, we find several genes related to ciliary basal body among our e-genes (*BBS2*; *CDKL5*; *CEP41*; *CFAP410*; *GAS8*), as well as another organelle also known to be implicated in metabolism, the lysosome (*CTSD*; *GALNS*; *HSPG2*; *SGSH*). As a matter of fact, our dataset includes several members of a gene family associated with Batten disease/ceroid lipofuscinosis, a lysosomal storage disorder (*CLN6*, *CLN7*/*MFSD8*, *CLN10*/*CTSD*).

Intersecting our e-gene list with several datasets providing richer annotations of cancient/modern genome differences (regions of positive selection; deserts of introgression; and reconstructed #d genome architectures) reveals the presence of a variant impacting *SLC12A5* associated with a signal of positive selection (Peyrégne et al., 2017). *SLC12A5* is a neuron specific membrane protein that is responsible for the maintenance of the low intracellular Cl-concentration in adult neurons. It plays a key role in GABAA and glycine signaling and is a point of convergence for multiple autism spectrum disorder and epilepsy risk gene products. Among these, *NETO2* is also associated with a signal of positive selection in *Homo sapiens*, and is also present in our e-gene list.

A few of our SNPs fall within regions with diverged 3D genome organization between *sapiens* and other hominins, as per the deep learning methods for inferring 3D genome organization from DNA sequence to Neanderthal, Denisovan, and diverse *sapiens* genomes in McArthur et al. (2025). Among them, QTLs with high NES are predicted to impact expression of *BBS2*, *HINT1*, and *RPL29*.

We identified 3 basal SNPs putatively re-introgressed in *sapiens* genomes via gene flow from Neanderthals (as per Kendall et al., 2025; Rong et al., 2023). One of them (rs77724956) corresponds to strongest sapiens-specific exonic splicing mutation in Rong et al. (2023). rs77724956 falls in the splice region of exon 12 of *FAAH*, a gene that encodes a catabolic enzyme for fatty acid amides such as anandamide, involved in pain sensation. Rong et al. (2023) report that the eQTLGen consortium, which provides whole-blood eQTLs based on 31,684 individuals across multiple ancestries, identifies rs77724956 as a strongly associated *FAAH* eQTL. We could not find information about this SNP in GTEx 10, but we note that it falls within a region of positive selection (Peyrégne et al., 2017).

The two other putatively re-introgressed (basal) SNPs (rs79625667; rs2306636) are eQTLs, and 1 of them is also a GWAS hit in a study about the genetic architecture of spatial electrical biomarkers for cardiac arrhythmia and relationship with cardiovascular disease. The SNP is associated with cardiovascular phenotypes in ClinVar.

Relatively few QTLs match GWAS hits in the EBI-GWAS database, as expected from the systematic differences in discovery of genetic effects on gene expression and complex traits (Mostafavi et al., 2023) (Table S13). Most matches correspond to height GWAS results. Most of these matches are cerebellum QTLs, which is consistent with the association between height and cerebellar phenotypes reported (for the UK biobank) in Wang et al. (2024).

Only one brain QTL result in this study is associated with a brain related GWAS (educational attainment). The tissues impacted (basal ganglia) are linked to expression of *PPP1R21*, a gene associated with an autosomal recessive intellectual disability syndrome, where a prominent and severe feature is the lack of speech or absent language skills (known to depend in part on basal ganglia structures).

## 3. Discussion

Our work interrogating the nature of derived variants whose frequencies diverge significantly among hominin genomes currently available fits within a growing attempt to harness information about variation withing contemporary human genomes (and associated phenotypes) in order to constrain our understanding of the functional impact of derived variants in *sapiens*, a situation where human conditions illuminate the human condition (Caporale et al., 2025). Unlike complementary attempts (Molz et al., 2024), which consider the effects of polymorphisms where both alleles tend to be relatively frequent, our work finds evidence that relatively rare minor alleles whose source can be traced back either far back in the primate lineage, or more exclusively to Neanderthal and Denisovan genomes, can point to ways in which *sapiens* may have come to be different from closely related but now-extinct hominins, especially when these variants are examined in the light of their associations with disease phenotypes.

Recurrent disease associations linked to specific polymorphic sites, coupled with supporting information about these SNPs regarding gene expression differences in specific tissues as well as cell type specificity of e-genes, add evidence for differences pertaining to the brain: depletion of brain-specific effects of introgressed alleles (a fact consistent with brain-specific expression profiles for genes found in large introgression deserts; Buisan et al., 2022; Vernot et al., 2016) and a marked impact of sapiens-derived alleles in specific tissues, most saliently the cerebellum. Gene Ontology enrichment results reinforce the existence of metabolic differences, but this specific study refines the set of distinguishing metabolic terms and points to an effect on astrocytes and other glial cells. While this effect remains to be tested further in a variety of experimental settings, it is in light of pronounced changes in human astrocytes compared to other extant primates (Hercher et al., 2024; Zintel et al., 2024; Rickelton et al., 2025; Ciuba et al., 2025). It would make eminent sense now that the important roles of astrocytes on neuronal networks, cognition and behavior are beginning to be recognized more explicitly (Barres, 2008; Eroglu, 2025; Kaul et al., 2024). Connections between astrocyte metabolism and the vascular system are emerging (D’Gama et al., 2025) and are potentially related to the white matter pathways recently highlighted in Molz et al. (2024) and Casas i Riu and Boeckx (2026), as well as the consistent enrichment of astrocytes among neuronal cell types reported in Andirkó et al. (2022). Isolated reports of strong effects in specific glial genes like *PHLPP1* (Gunz et al., 2019), *DLAT* (Alagöz et al., 2024) or PC (Prüfer et al., 2014), alongside metabolic differences (lactate, purine biosynthesis) that likely impact astrocytes (*ADSL*, *SLC16A1*), deserve further investigation in light of our results. This will require developing experimental models (e.g., organoids) that still rarely feature in standard toolkits (but see Sloan et al., 2017). At the moment, a rewiring leading to a more extensive use of the glycolytic pathway in *sapiens* emerges as a hypothesis worth exploring, especially considering early literature linking aerobic glycolysis (in neurons) to neoteny (Goyal et al., 2024; Bufill et al., 2011; Somel et al., 2009).

In addition to glia, our results encourage future work on lysosomes (reinforcing the relevance of autophagy pathways for brain development and function; Leonardi et al., 2025; Skaros et al., 2025; Lewerissa et al., 2024) and cilia. Here, too, our focus on differences among hominins highlights themes already identified in comparative studies focusing on living primates (Dumas et al., 2021). This more general trend separating *Homo* from *Pan* may have been accentuated in the *sapiens* lineage, where cilia-related differences and Joubert-syndrome-like phenotypes have occasionally been referenced (Plomp et al., 2025; Funato et al., 2024; Takagi & Kondo, 2026).

While each SNP result could be relevant, and deserves further exploration, we find it noteworthy that quite a few genes predicted to be associated with the SNPs in our study (several members of CACNA1 gene family, *MAOA*, *GRIN2A*, *FAAH*, *SLITRK1*, FMR1), and indeed several e-genes (such as *CDKL5*, *NETO2* and *SLC12A5*, but also *KCNN3* and *CHRM5*), are related to neuronal transmission and its regulation, critical to explain hypothesized differences in the modulation of the stress response among hominins (O’Rourke & Boeckx, 2020; O’Rourke et al., 2021).

It bears repeating that a full characterization of the differences observed here require additional experimental validation. Among other things, this will require taking on board the recognition that a single SNP is likely to impact multiple genes (Lawrence et al., 2025), making investigations more difficult. Likewise, the difference in differential gene expression detection between the basal and derived datasets points to a frequency effect that may limit the insights one can gain from large GWAS results, whose robustness is often frequency-dependent. This makes the detecting of convergence between common and rare variants more difficult (Spence et al., 2026), requiring a pathway-centric approach instead of a variant-or even gene-centric one (Wright et al., 2025).

## Methods

### Original data

We extracted the HF variants from the supplementary materials in Kuhlwilm and Boeckx’s (2019) catalog, which used three high-coverage genotypes for the extinct human population (one Denisovan, Denisova 3 and two Neanderthals, Vindija 33.19 and Denisova 5, also known as the Altai Neandertal) and the human genome hg19 as reference for the living human population. Relevant variants were identified in positions where more than 90% of present-day humans carry a (*sapiens*-)derived allele while at least the Denisovan and one Neanderthal carry the ancestral allele. Published data on multiple genome alignments (Paten et al., 2008) was used to infer ancestral alleles, and in positions where this information was missing, the authors used the macaque reference allele, *rheMac3* (Yan et al., 2011). Moreover Kuhlwilm and Boeckx (2019) determined allele frequencies using the dbSNP database (Sherry et al., 2001). Here minor allele frequencies were derived from the human derived allele frequency (human_DAF) values in the basal and derived variant catalogues. For basal variants, we calculated MAF as 1 − human_DAF, whereas for derived variants, human_DAF was used directly. Mean MAF was calculated for each group, and the two groups were compared using a two-sided Welch’s two-sample t-test.

Additionally, we further filtered basal and derived HF variants by intersecting them with variants that are identified in individual patient samples as reported by the Clinical Variation (ClinVar) database (https://www.ncbi.nlm.nih.gov/clinvar/). We plotted our final list of close to 2000 loci on the karyotype using the karyoploteR package in R to visualize the localisation of the retained variants after filtering.

Upon ClinVar-based filtering and obtaining a list further divided into the two basal and derived subsets, we first aggregated variant metadata to provide genomic/descriptive context, before annotating them functionally using various databases, to finally perform downstream statistical analyses.

### Disease Associations

Firstly, each variant had a ClinVar entry specifying the disease phenotype(s) with which it has been reported to be associated. Because the terminology in the ClinVar reports is varied, we normalized the disease labels by converging some diagnoses to detect patterns of concentration. We further excluded the highly common term “Inborn genetic disease” as it was too generic to be interpreted.

### Gene Associations; gene length

Then, we mapped variants to relevant genes using the GREAT (Genomic Regions Enrichment of Annotations Tool) platform (https://great.stanford.edu/). This tool defines regulatory domains by linking non-coding DNA sequences to nearby genes, considering both their immediate proximity and extended regulatory domains reaching up to 1 Mb. Moreover, from Kuhlwilm and Boeckx 2019, we extracted the genomic length of all genes reported to be associated with our variants. To assess whether recurrent disease associations in our dataset could be influenced by gene length, we examined the genomic lengths of genes associated with the ten most frequently represented disease categories in the basal and (N-D)-derived variant sets. Within the top-ten disease subset, each distinct gene was counted only once, yielding 119 basal and 146 derived genes. One gene from each subset had no corresponding length entry and was excluded, leaving 118 basal and 145 derived genes. For each set, an equally sized random control sample was selected without replacement from the corresponding eligible gene pool, excluding disease-associated genes. Gene lengths were compared between disease-associated and control genes using two-sided Welch’s two-sample t-tests (random seed = 123; Table S3).

### GTEx

We introduced all variants into the Genotype-Tissue Expression (GTEx) V10 portal, which reports variants identified as expressive/splicing Quantitative Trait Loci (e-/sQTLs) in specific tissues. We discarded those returning null results and generated a table with associated gene(s), tissue-specific effects, and normalized effect size (NES).

Moreover, we examined whether the distribution of tissues in our dataset with identified QTLs deviated from the proportions expected given GTEx sample sizes for each tissue. Fisher’s exact tests were performed, with false discovery rate and Bonferroni correction applied to control for false positives. To identify tissues contributing most strongly to observed deviations, we inspected standardized residuals from chi-squared tests; however, because some GTEx tissues contained fewer than five RNA samples, we used the residuals only and did not interpret chi-squared p-values.

To test whether brain-associated SNPs were overrepresented in either group, we performed a permutation analysis in which group labels (basal vs. derived) were randomly reassigned across SNPs while preserving brain/non-brain status. This procedure was repeated 10,000 times to generate a null distribution of brain SNP counts in the derived group, from which a p-value was derived.

### GnomAD

We further investigated potential functional implications of our variants. To characterize genetic constraint, defined as the tolerance of genes or genomic regions to genetic variation, we assessed both gene-level and genomic-region constraint using metrics from gnomAD. The proportion of coding and non-coding variants was determined from the Ensembl VEP biotype and mutation consequence annotations. Although all variants were associated with protein-coding genes, their predicted consequences included both coding and non-coding variants. Gene-level constraint was used to assess the tolerance of the associated protein-coding gene to variation, while Gnocchi (Genomic Constraint Score) was used to assess constraint of the genomic region surrounding each variant.

To annotate the variants with gene constraint scores, VEP-annotated variants were first matched to transcript identifiers in the gnomAD v4.1.1 constraint dataset. Only MANE Select and MANE Plus Clinical transcripts were retained as transcript references. Constraint scores were obtained according to the predicted consequence of each variant. For missense and synonymous variants, the corresponding Z-scores were used, with higher positive values indicating greater intolerance to variation and therefore greater constraint. For pLoF variants, LOEUF was used, defined as the upper bound of the 90% confidence interval for the observed/expected ratio of high-confidence pLoF variants, with lower values indicating greater constraint. Extreme synonymous Z-scores were considered indicative of potential data quality issues. For each variant, the constraint score corresponding to its predicted consequence was assigned at the gene level. Thus, synonymous variants were assigned the synonymous constraint score (syn.z_score), missense variants were assigned the missense constraint score (mis.z_score), and pLoF variants were assigned LOEUF. A gene could therefore be included in more than one consequence class, with the appropriate constraint score used for each class. Duplicate gene-score combinations were counted only once. Basal and derived groups were compared separately within each consequence class using two-sided Wilcoxon rank-sum tests. Mean and median constraint scores were also calculated for each group.

Genomic constraint was assessed using Gnocchi, a genome-wide constraint score that measures the depletion of observed genetic variation relative to the expected level of variation within 1-kb genomic windows. Higher positive Gnocchi scores indicate greater genomic constraint. Each SNP was assigned a Gnocchi score based on its genomic position and the nearest 1-kb genomic window. SNPs with missing scores were excluded from the analysis (basal n = 658, derived n = 932).

To compare genomic constraint between basal and derived variants, Gnocchi (constraint_z) score distributions were assessed for normality using Shapiro–Wilk tests. Both groups significantly deviated from normality (p < 0.001). Differences between the groups were assessed using a two-sided Wilcoxon rank-sum test. A Welch two-sample t-test was additionally used to compare mean Gnocchi scores. Mean, median and standard deviation were calculated for each group, and Cohen’s d was calculated to quantify the effect size of the difference in mean scores.

### Mutation type and mutation direction

The predicted mutation consequence of each variant was obtained using the Ensembl Variant Effect Predictor (VEP). VEP annotations were restricted to entries matching the same rsID and alternate allele as our variants, ensuring that the predicted consequence corresponded to the observed variant. Only MANE Select and MANE Plus Clinical transcripts were retained as transcript references. These annotations provided the predicted mutation consequence, biotype and VEP impact category also used in subsequent analyses.

The VEP mutation consequences were subsequently used to characterize the distribution of variant types between basal and derived variants. Because individual rsIDs could have multiple annotations, the statistical analyses accounted for this dependency. An initial permutation test was used to test whether the overall distribution of mutation consequences differed between basal and derived variants. To assess the direction of this difference, VEP consequences were grouped according to their predicted impact (MODIFIER, LOW, MODERATE and HIGH) and analysed using an ordinal logistic regression model. This showed whether derived variants were more likely to have higher predicted-impact consequences.

### CADD

To assess the predicted functional impact of individual variants, we evaluated their CADD scores. CADD integrates more than 60 genomic annotations to estimate the predicted deleteriousness or functional impact of individual variants, with higher scores indicating greater predicted impact. PHRED-scaled CADD scores rank variants relative to approximately 9 billion possible human single-nucleotide variants, with a score of 10 corresponding approximately to the top 10% and 20 to the top 1%.

We extracted PHRED scores for the basal and derived variants and compared their distributions. Of the 711 basal variants, 431 (60.6%) had an available CADD PHRED score, compared with 505 of 957 (52.8%) derived variants. Summary statistics were calculated for each group, and pairwise Welch two-sample t-tests were performed across groups. Bonferroni correction was applied to account for multiple comparisons, and Cohen’s d was calculated to estimate effect size. Pre-Bonferroni correction, the only significant comparison was between derived eQTL and derived sQTL variants. Comparisons between brain-related and non-brain-related QTLs were not possible because only one brain-related QTL variant had an available CADD PHRED score.

### Dates of Emergence and additional SNP information

Moreover, we made use, when possible, of available evolutionary information for each SNP by indicating the date of emergence. Emergence dates for the variants were obtained thanks to the Genealogical Estimation of Variant Age (GEVA) tool, which uses a hidden Markov model approach to infer the location of ancestral haplotypes relative to a given variant, as per Andirkó et al. (2022). Coordinates for regions showing signals of positive selection were obtained from the supplementary files provided by Peyrégne et al. (2017) and desert of introgression coordinates were extracted from Chen et al. (2020). We also checked the geographical distribution of the N-D derived variants using the 1000genome platform (https://www.internationalgenome.org/).

### Enrichments

Following gene association, we performed enrichment (gene ontology pathway) analyses to explore whether genes linked to the variant sets were over-represented in specific biological processes (BP), molecular functions (MF), and cellular components (CC), using primarily the topGO package provided by R, and secondarily verifying result validity with gProfiler in order to have double source confirmation. Enrichment analyses were conducted on the following subsets: all basal variants; all derived variants; all basal variants whose associated genes appear on gTEX and within those all that are expressed in the brain, and in the cerebellum/cerebellar hemisphere; all derived variants; all derived variants whose associated genes appear on gTEX and within those all that are expressed in the brain, and in the cerebellum/cerebellar hemisphere. Because enrichment analyses yield large datasets which are difficult to interpret, it is common procedure to apply various reduction techniques to obtain most informative terms. We used the GOSemSim package provided by R which groups the enrichment into a chosen number of clusters. We chose 15 clusters. Then, we selected one representative term for each cluster based on the criterion that the term contained at least two overlapping genes; and among those, the term with the lowest weight01Fisher value was selected.

Additionally, we ran a separate set of enrichment analyses on the basal and derived variants which had associated genes that appeared in GTEx and had QTLs which were expressed in the brain, using the brain-specific functional enrichment platform SYNGO which provides synaptic specific gene ontologies.

### Cell Type

Upon identifying enriched biological processes and putative regulatory mechanisms through transcription factor binding analyses, we subsequently annotated the brain-related variants with cell-type specific information to contextualize these findings within specific cellular populations and to refine the biological interpretation of the observed functional patterns. For this cell type annotation, we used the CELLxGENE expression platform (https://cellxgene.cziscience.com/gene-expression) and extracted for each gene the provided expression values over different cell types (derived from single-cell RNA sequencing data), along with the provided Number of Cells Expressing Genes. We subsequently ranked cell type frequency based on the number of genes from our list, whose scaled expression was highest in each cell type. We performed this annotation for basal eQTLs; basal sQTLs; derived eQTLs and derived sQTLs.

### SFARI genes

To further investigate the potential functional relevance of variants associated with gene expression or splicing, we annotated the associated genes using the Simons Foundation Autism Research Initiative (SFARI) database (https://gene.sfari.org/). For genes present in SFARI, we extracted their genetic category and available gene-level annotations, including whether the association was syndromic and the number of supporting reports. These annotations were used to compare the presence and characteristics of SFARI-associated genes among basal and derived eQTLs and sQTLs.

### GWAS catalog

We further interrogated the potential pathogenicity of our variants associated with gene expression or splicing by intersecting them with variants reported in the NHGRI-EBI GWAS Catalog (https://www.ebi.ac.uk/gwas/), a curated catalogue of genetic variants previously reported in genome-wide association studies (GWAS) and associated with diseases or complex traits.

## Acknowledgments

We thank Juan Moriano for assistance and valuable feedback at the early stage of this project, and Judit Casas i Riu for valuable comments on the draft.

## Author contributions

Conceptualization: C. B.; methodology: C. B., E.M.Y.; data curation: E.M.Y.; formal analysis: E.M.Y., G.B., Y.Y., C.B.; visualization: E.M.Y., G.B..; writing – original draft: C. B., E.M.Y.; writing – review and editing: E.M.Y., G.B., Y.Y., C.B; supervision: C. B.; funding acquisition: C. B.

## References

Alagöz, G., Eising, E., Mekki, Y., Bignardi, G., Fontanillas, P., 23andMe Research Team, Aslibekyan, S., Auton, A., Babalola, E., Bell, R. K., Bielenberg, J., Bowes, J., Bryc, K., Chaudhary, N. S., Coker, D., Das, S., DelloRusso, E., Elson, S. L., Eriksson, N., … Gordon, R. L. (2024). The shared genetic architecture and evolution of human language and musical rhythm. Nature Human Behaviour, 9(2), 376–390. 10.1038/s41562-024-02051-y

Albers, P. K., & McVean, G. (2020). Dating genomic variants and shared ancestry in population-scale sequencing data. PLoS Biology, 18(1), e3000586. 10.1371/journal.pbio.3000586

Andirkó, A., & Boeckx, C. (2022). Brain region-specific effects of nearly fixed *sapiens*-derived alleles. BMC Genomic Data, 23(1), 36. 10.1186/s12863-022-01048-8

Andirkó, A., Moriano, J., Vitriolo, A., Kuhlwilm, M., Testa, G., & Boeckx (2022). Temporal mapping of derived high-frequency gene variants supports the mosaic nature of the evolution of *Homo sapiens*. Scientific Reports, 12, 9937. 10.1038/s41598-022-13589-0

Barres, B. A. (2008). The mystery and magic of glia: a perspective on their roles in health and disease. Neuron, 60(3), 430–440. 10.1016/j.neuron.2008.10.013

Brand, C. M., Colbran, L. L., & Capra, J. A. (2022). Predicting archaic hominin phenotypes from genomic data. Annual Review of Genomics and Human Genetics, 23, 591–612. 10.1146/annurev-genom-111521-121903

Bufill, E., Agustí, J., & Blesa, R. (2011). Human neoteny revisited: The case of synaptic plasticity. American Journal of Human Biology, 23(6), 729–739. 10.1002/ajhb.21225

Buisan, R., Moriano, J., Andirkó, A., & Boeckx, C. (2022). A brain region-specific expression profile for genes within large introgression deserts and under positive selection in *Homo sapiens*. Frontiers in Cell and Developmental Biology, 10, 824740. 10.3389/fcell.2022.824740

Caporale, N., Leonardi, O., Villa, C. E., Vitriolo, A., Boeckx, C., & Testa, G. (2025). Tile by tile: capturing the evolutionary mosaic of human conditions. Current Opinion in Genetics & Development, 90, 102297. 10.1016/j.gde.2024.102297

Casas i Riu, J. & Boeckx, C. (2026). Contextualizing the polygenic basis of human brain shape. bioRxiv 2026.10.01.754889. 10.64898/2026.10.01.754889

Chen, L., Wolf, A. B., Fu, W., Li, L., & Akey, J. M. (2020). Identifying and interpreting apparent Neanderthal ancestry in African individuals. Cell, 180(4), 677–687.e16. 10.1016/j.cell.2020.01.012

Ciuba, K., Piotrowska, A., Chaudhury, D., Dehingia, B., Duński, E., Behr, R., Soroczyńska, K., Czystowska-Kuźmicz, M., Abbas, M., Bulanda, E., Gawlik-Zawiślak, S., Pietrzak, S., Figiel, I., Włodarczyk, J., Verkhratsky, A., Niedbała, M., Kaspera, W., Wypych, T., Wilczyński, B., & Pękowska, A. (2025). Molecular signature of primate astrocytes reveals pathways and regulatory changes contributing to human brain evolution. Cell Stem Cell, 32(3), 426–444.e14. 10.1016/j.stem.2024.12.011

D’Gama, P. P., Jeong, I., Nygård, A. M., Jamali, A., Yaksi, E., & Jurisch-Yaksi, N. (2025). Motile cilia modulate neuronal and astroglial activity in the zebrafish larval brain. Cell Reports, 44(1), 115195. 10.1016/j.celrep.2024.115195

Ding, Y., Hou, K., Xu, Z., Pimplaskar, A., Petter, E., Boulier, K., Privé, F., Vilhjálmsson, B. J., Olde Loohuis, L. M., & Pasaniuc, B. (2023). Polygenic scoring accuracy varies across the genetic ancestry continuum. Nature, 618(7966), 774–781. 10.1038/s41586-023-06079-4

Dumas, G., Malesys, S., & Bourgeron, T. (2021). Systematic detection of brain protein-coding genes under positive selection during primate evolution and their roles in cognition. Genome Research, 31(3), 484–496. 10.1101/gr.262113.120

Eroglu C. (2025). Astrocytes, hidden puppet masters of the brain. Science, 388(6748), 705–706. 10.1126/science.adx7102

Funato, N., Heliövaara, A., & Boeckx, C. (2024). A regulatory variant impacting TBX1 expression contributes to basicranial morphology in *Homo sapiens*. The American Journal of Human Genetics, 111(5), 939–953. 10.1016/j.ajhg.2024.03.012

Gokcumen, O. (2020). Archaic hominin introgression into modern human genomes. American Journal of Physical Anthropology, 171(Suppl. 70), 60–73. 10.1002/ajpa.23951

Goyal, M. S., Hawrylycz, M., Miller, J. A., Snyder, A. Z., & Raichle, M. E. (2014). Aerobic glycolysis in the human brain is associated with development and neotenous gene expression. Cell Metabolism, 19(1), 49–57. 10.1016/j.cmet.2013.11.020

Gunz, P., Tilot, A. K., Wittfeld, K., Teumer, A., Shapland, C. Y., van Erp, T. G. M., Dannemann, M., Vernot, B., Neubauer, S., Guadalupe, T., Fernández, G., Brunner, H. G., Enard, W., Fallon, J., Hosten, N., Völker, U., Profico, A., Di Vincenzo, F., Manzi, G., Kelso, J., … Fisher, S. E. (2019). Neandertal introgression sheds light on modern human endocranial globularity. Current Biology, 29(1), 120–127.e5. 10.1016/j.cub.2018.10.065

Hercher, C., Ellerbeck, K., Toutée, L., Ye, X., Mpai, R., Davoli, M. A., Watt, A. J., Murai, K. K., Turecki, G., & Mechawar, N. (2024). Human-specific features of cerebellar astrocytes and Purkinje cells: An anatomical comparison with mice and macaques. bioRxiv, 2024.08.13.607849. 10.1101/2024.08.13.607849

Huang, X., Hackl, J., & Kuhlwilm, M. (2025). Decoding genomic landscapes of introgression. Trends in Genetics, 41(12), 1096–1108. 10.1016/j.tig.2025.07.001

Kaul, D., Curry, A. R., Gerstner, N., Fröhlich, A. S., Krontira, A. C., Kyrousi, C., Smith, C. C., Sutherland, G. T., Dottori, M., Ziller, M. J., Chatzinakos, C., Daskalakis, N. P., Cappello, S., Czamara, D., Binder, E. B., Cruceanu, C., Knauer-Arloth, J., Mechawar, N., Schwab, S. G., … Matosin, N. (2024). Astrocytic glutamate regulation is shaped by adversity and glucocorticoid signalling. bioRxiv, 2024.06.24.600362. 10.1101/2024.06.24.600362

Kendall, C., Nooranikhojasteh, A., Parra, E. J., Schillaci, M. A., & Viola, B. (2025). Archaic adaptive introgression in modern human reproductive genes. Communications Biology, 8(1), 1365. 10.1038/s42003-025-08682-9

Kuhlwilm, M., & Boeckx, C. (2019). A catalog of single nucleotide changes distinguishing modern humans from archaic hominins. Scientific Reports, 9, 8463. 10.1038/s41598-019-44877-x

Lawrence, K., Gjorgjieva, T., & Montgomery, S. B. (2025). Focus on single gene effects limits discovery and interpretation of complex trait-associated variants. bioRxiv, 2025.06.06.658175. 10.1101/2025.06.06.658175

Leonardi, O., Lewerissa, E., Aprile, D., Barakat, T. S., Culotta, L., Deng, R., Haazen, L., Linda, K., Majoie, M., Mordelt, A., Ortale, A., Oudakker, A., Prazzoli, F., Puvogel, S., Scheefhals, N., Schelhaas, H. J., Schoenmaker, C., Van Hughte, E., Van Bokhoven, H., … Kasri, N. N. (2025). Human-specific regulation of CHD2 controls the tempo of cortical development through lysosomal homeostasis. bioRxiv, 2025.01.21.634145. 10.1101/2025.01.21.634145

Lewerissa, E. I., Nadif Kasri, N., & Linda, K. (2024). Epigenetic regulation of autophagy-related genes: Implications for neurodevelopmental disorders. Autophagy, 20(1), 15–28. 10.1080/15548627.2023.2250217

Mafessoni, F., Grote, S., de Filippo, C., Slon, V., Kolobova, K. A., Viola, B., Markin, S. V., Chintalapati, M., Peyrégne, S., Skov, L., Skoglund, P., Krivoshapkin, A. I., Derevianko, A. P., Meyer, M., Kelso, J., Peter, B., Prüfer, K., & Pääbo, S. (2020). A high-coverage Neandertal genome from Chagyrskaya Cave. Proceedings of the National Academy of Sciences of the United States of America, 117(26), 15132–15136. 10.1073/pnas.2004944117

McArthur, E., Rinker, D. C., Cheng, Y., Wang, Q., Wang, J., Gilbertson, E. N., Fudenberg, G., Pittman, M., Keough, K., Yue, F., Pollard, K. S., & Capra, J. A. (2025). Reconstructing the 3D genome organization of Neanderthals reveals that chromatin folding shaped phenotypic and sequence divergence. bioRxiv, 2022.02.07.479462. 10.1101/2022.02.07.479462

McCoy, R. C., Wakefield, J., & Akey, J. M. (2017). Impacts of Neanderthal-introgressed sequences on the landscape of human gene expression. Cell, 168(5), 916–927.e12. 10.1016/j.cell.2017.01.038

Molz, B., Eising, E., Alagöz, G., Schijven, D., Francks, C., Gunz, P., & Fisher, S. E. (2024). Imaging genomics reveals genetic architecture of the globular human braincase. bioRxiv, 2024.03.20.585712. 10.1101/2024.03.20.585712

Mostafavi, H., Spence, J. P., Naqvi, S., & Pritchard, J. K. (2023). Systematic differences in discovery of genetic effects on gene expression and complex traits. Nature Genetics, 55(11), 1866–1875. 10.1038/s41588-023-01529-1

Neubauer, S., Hublin, J. J., & Gunz, P. (2018). The evolution of modern human brain shape. Science Advances, 4(1), eaao5961. 10.1126/sciadv.aao5961

Ongaro, L., & Huerta-Sanchez, E. (2024). A history of multiple Denisovan introgression events in modern humans. Nature Genetics, 56, 2612–2622. 10.1038/s41588-024-01960-y

O’Rourke, T., & Boeckx, C. (2020). Glutamate receptors in domestication and modern human evolution. Neuroscience & Biobehavioral Reviews, 108, 341–357. 10.1016/j.neubiorev.2019.10.004

O’Rourke, T., Martins, P. T., Asano, R., Tachibana, R. O., Okanoya, K., & Boeckx, C. (2021). Capturing the effects of domestication on vocal learning complexity. Trends in Cognitive Sciences, 25(6), 462–474. 10.1016/j.tics.2021.03.007

Pääbo, S. (2014). The human condition—a molecular approach. Cell, 157(1), 216–226. 10.1016/j.cell.2013.12.036

Paten, B., Herrero, J., Fitzgerald, S., Beal, K., Flicek, P., Holmes, I., & Birney, E. (2008). Genome-wide nucleotide-level mammalian ancestor reconstruction. Genome Research, 18(11), 1829– 1843. 10.1101/gr.076521.108

Peyrégne, S., Boyle, M. J., Dannemann, M., & Prüfer, K. (2017). Detecting ancient positive selection in humans using extended lineage sorting. Genome Research, 27(9), 1563–1572. 10.1101/gr.219493.116

Plomp, K., Lewis, D., Buck, L., Bukhari, S., Rae, T., Gnanalingham, K., & Collard, M. (2025). A test of the Archaic *Homo* Introgression Hypothesis for the Chiari malformation type I. Evolution, Medicine, and Public Health, 13(1), 154–166. 10.1093/emph/eoaf009

Prüfer, K., Racimo, F., Patterson, N., Jay, F., Sankararaman, S., Sawyer, S., Heinze, A., Renaud, G., Sudmant, P. H., de Filippo, C., Li, H., Mallick, S., Dannemann, M., Fu, Q., Kircher, M., Kuhlwilm, M., Lachmann, M., Meyer, M., Ongyerth, M., Siebauer, M., … Pääbo, S. (2014). The complete genome sequence of a Neanderthal from the Altai Mountains. Nature, 505(7481), 43–49. 10.1038/nature12886

Rickelton, K., Babbitt, C. C., & Babbitt, C. (2025). Exploring the expanded role of astrocytes in primate brain evolution via changes in gene expression. *Brain*, Behavior and Evolution, 100(3), 200–208. 10.1159/000544004

Rinker, D. C., Simonti, C. N., McArthur, E., Shaw, D., Hodges, E., & Capra, J. A. (2020). Neanderthal introgression reintroduced functional ancestral alleles lost in Eurasian populations. Nature Ecology & Evolution, 4(10), 1332–1341. 10.1038/s41559-020-1261-z

Rong, S., Neil, C. R., Welch, A., Duan, C., Maguire, S., Meremikwu, I. C., Meyerson, M., Evans, B. J., & Fairbrother, W. G. (2023). Large-scale functional screen identifies genetic variants with splicing effects in modern and archaic humans. Proceedings of the National Academy of Sciences of the United States of America, 120(21), e2218308120. 10.1073/pnas.2218308120

Sherry, S. T., Ward, M. H., Kholodov, M., Baker, J., Phan, L., Smigielski, E. M., & Sirotkin, K. (2001). dbSNP: the NCBI database of genetic variation. Nucleic Acids Research, 29(1), 308–311. 10.1093/nar/29.1.308

Siletti, K., Hodge, R., Mossi Albiach, A., Lee, K. W., Ding, S. L., Hu, L., Lönnerberg, P., Bakken, T., Casper, T., Clark, M., Dee, N., Gloe, J., Hirschstein, D., Shapovalova, N. V., Keene, C. D., Nyhus, J., Tung, H., Yanny, A. M., Arenas, E., Lein, E. S., … Linnarsson, S. (2023). Transcriptomic diversity of cell types across the adult human brain. Science, 382(6667), eadd7046. 10.1126/science.add7046

Skaros, A., Vitriolo, A., Leonardi, O., Finazzi, V., Pereira, M. F., Prazzoli, F., Trattaro, S., Moriano, J., Capocefalo, D., Villa, C. E., Boettcher, M., Boeckx, C., & Testa, G. (2025). Regulatory logic of human cortex evolution by combinatorial perturbations. bioRxiv, 2025.04.28.651083. 10.1101/2025.04.28.651083

Skov, L., Coll Macià, M., Sveinbjörnsson, G., Mafessoni, F., Lucotte, E. A., Einarsdóttir, M. S., Jonsson, H., Halldorsson, B., Gudbjartsson, D. F., Helgason, A., Schierup, M. H., & Stefansson, K. (2020). The nature of Neanderthal introgression revealed by 27,566 Icelandic genomes. Nature, 582(7810), 78–83. 10.1038/s41586-020-2225-9

Sloan, S. A., Darmanis, S., Huber, N., Khan, T. A., Birey, F., Caneda, C., Reimer, R., Quake, S. R., Barres, B. A., & Paşca, S. P. (2017). Human astrocyte maturation captured in 3D cerebral cortical spheroids derived from pluripotent stem cells. Neuron, 95(4), 779–790.e6. 10.1016/j.neuron.2017.07.035

Somel, M., Franz, H., Yan, Z., Lorenc, A., Guo, S., Giger, T., Kelso, J., Nickel, B., Dannemann, M., Bahn, S., Webster, M. J., Weickert, C. S., Lachmann, M., Pääbo, S., & Khaitovich, P. (2009). Transcriptional neoteny in the human brain. Proceedings of the National Academy of Sciences, 106(14), 5743–5748. 10.1073/pnas.0900544106

Spence, J. P., Mostafavi, H., Ota, M., Milind, N., Gjorgjieva, T., Smith, C. J., Simons, Y. B., Sella, G., & Pritchard, J. K. (2026). Specificity, length and luck drive gene rankings in association studies. Nature, 649(8098), 918–925. 10.1038/s41586-025-09703-7

Takagi, K., & Kondo, O. (2026). Postnatal interaction of size and shape in the human endocranium and brain structures. Journal of Anatomy, 248(3), 395–411. 10.1111/joa.70040

Vernot, B., Tucci, S., Kelso, J., Schraiber, J. G., Wolf, A. B., Gittelman, R. M., Dannemann, M., Grote, S., McCoy, R. C., Norton, H., Scheinfeldt, L. B., Merriwether, D. A., Koki, G., Friedlaender, J. S., Wakefield, J., Pääbo, S., & Akey, J. M. (2016). Excavating Neandertal and Denisovan DNA from the genomes of Melanesian individuals. Science, 352(6282), 235–239. 10.1126/science.aad9416

Wang, Z., Diedrichsen, J., Saltoun, K., Steele, C., Arnold-Anteraper, S. R., Yeo, B. T. T., Schmahmann, J. D., & Bzdok, D. (2024). Structural covariation between cerebellum and neocortex intrinsic structural covariation links cerebellum subregions to the cerebral cortex. Journal of Neurophysiology, 132(3), 849–869. 10.1152/jn.00164.2024

Weiss, C. V., Harshman, L., Inoue, F., Fraser, H. B., Petrov, D. A., Ahituv, N., & Gokhman, D. (2021). The *cis*-regulatory effects of modern human-specific variants. eLife, 10, e63713. 10.7554/eLife.63713

Wen, C., Margolis, M., Dai, R., Zhang, P., Przytycki, P. F., Vo, D. D., Bhattacharya, A., Matoba, N., Tang, M., Jiao, C., Kim, M., Tsai, E., Hoh, C., Aygün, N., Walker, R. L., Chatzinakos, C., Clarke, D., Pratt, H., PsychENCODE Consortium, Peters, M. A., … Gandal, M. J. (2024). Cross-ancestry atlas of gene, isoform, and splicing regulation in the developing human brain. Science, 384(6698), eadh0829. 10.1126/science.adh0829

Wright, S. N., Yang, J., & Ideker, T. (2025). Common and rare genetic variants show network convergence for a majority of human traits. medRxiv, 2025.06.27.25330419. 10.1101/2025.06.27.25330419

Yan, G., Zhang, G., Fang, X., Zhang, Y., Li, C., Ling, F., Cooper, D. N., Li, Q., Li, Y., Van Gool, A. J., Du, H., Chen, J., Chen, R., Zhang, P., Huang, Z., Thompson, J. R., Meng, Y., Bai, Y., Wang, J., … Wang, J. (2011). Genome sequencing and comparison of two nonhuman primate animal models, the cynomolgus and Chinese rhesus macaques. Nature Biotechnology, 29(11), 1019– 1023. 10.1038/nbt.1992

Zeberg, H., Jakobsson, M., & Pääbo, S. (2024). The genetic changes that shaped Neandertals, Denisovans, and modern humans. Cell, 187, 1047–1058. 10.1016/j.cell.2023.12.029

Zintel, T. M., Pizzollo, J., Claypool, C. G., & Babbitt, C. C. (2024). Astrocytes drive divergent metabolic gene expression in humans and chimpanzees. Genome Biology and Evolution, 16(1), evad239. 10.1093/gbe/evad239

Zylka, M. J., Simon, J. M., & Philpot, B. D. (2015). Gene length matters in neurons. Neuron, 86(2), 353–355. 10.1016/j.neuron.2015.03.059

