## Supplementary material for "From ancestral patterns of variation to the human condition: Leveraging disease phenotype associations and gene expression changes": Suppl. Material: Legends_suppl_material.docx

**1. Variants dataset**

1_variants_dataset.xlsx

basal

derived

**2. Disease associations dataset**

2_disease_associations_dataset.xlsx

basal

basal null results

derived

derived null results

**3. Gene associations**

3_gene_associations.xlsx

basal

derived

basal genomic lengths

derived genomic lengths

basal random genomic lengths

derived random genomic lengths

genomic lengths test summary

**4. GTEx**

4_GTEx.xlsx

basal eqtl nes

basal sqtl nes

derived eqtl nes

derived sqtl nes

GTex sampling

basal brain eqtl residuals

basal brain sqtl residuals

derived brain sqtl residuals

**5. gnomAD**

5_gnomAD.xlsx

genomic constraint score

gene constraint score basal

gene constraint score derived

gene constraint test summary

genomic constraint test summary

**6. Mutation type and direction**

6_mutation_type_and_direction.xlsx

basal vep mane

derived vep mane

**7. CADD**

7_CADD.xlsx

basal

derived

cadd_pairwise_test_results

**8. SynGO enrichments**

8_SynGO_enrichments.xlsx

basal

derived

**9. Dates of emergence**

9_dates_of_emergence.xlsx

basal

brain

positive selection

**10. topGO enrichments**

10_topGO_enrichments.xlsx

GTEx brain (BP, MF, CC)

Basal brain (BP, MF, CC)

Basal eQTL brain (BP, MF, CC)

Basal sQTL brain (BP, MF, CC)

Derived brain (BP, MF, CC)

Derived eQTL brain (BP, MF, CC)

Derived sQTL brain (BP, MF, CC)

Cerebellum and hemisphere (BP, MF, CC)

Fetal-QTL (BP, MF, CC)

**11. Cell type**

11_cell_type.xlsx

basal eqtl

basal sqtl

derived eqtl

derived sqtl

confidence genes

cerebellar genes

**12. SFARI**

12_SFARI.xlsx

basal eqtl

basal sqtl

derived eqtl

derived sqtl

**13. GWAS**

13_GWAS.xlsx

basal eqtl

basal sqtl

derived eqtl

**14. GTEx v8 vs v10**

14_GTEx_v8_vs_v10.xlsx

GTEx v8 vs v10
